# Characterization of brown adipose tissue thermogenesis in the naked mole-rat (*Heterocephalus glaber*), a poikilothermic mammal

**DOI:** 10.1101/2020.04.28.062737

**Authors:** Yuki Oiwa, Kaori Oka, Hironobu Yasui, Kei Higashikawa, Hidemasa Bono, Yoshimi Kawamura, Shingo Miyawaki, Akiyuki Watarai, Takefumi Kikusui, Atsushi Shimizu, Hideyuki Okano, Yuji Kuge, Kazuhiro Kimura, Yuko Okamatsu-Ogura, Kyoko Miura

## Abstract

The naked mole-rat (NMR) is a poikilothermic mammal that forms eusocial colonies consisting of one breeding queen, several breeding kings, and subordinates. Despite their poikilothermic feature, NMRs possess brown adipose tissue (BAT), which in homeothermic mammals induces thermogenesis in cold environments. However, NMR-BAT thermogenic potential is controversial, and its physiological roles are unknown. Here, we show that NMR-BAT has beta-3 adrenergic receptor (ADRB3)-dependent thermogenic potential, which contributes to thermogenesis in the isolated queen in non-cold environments. NMR-BAT expressed several brown adipocyte marker genes and showed noradrenaline-dependent thermogenic activity *in vitro* and *in vivo*. Although our ADRB3 inhibition experiments revealed that NMR-BAT thermogenesis slightly delays the decrease in body temperature in a cold environment, it was insufficient to maintain the body temperatures of the NMRs. In a non-cold environment, NMRs are known to increase their body temperature by a heat-sharing behavior. Interestingly, we found that the body temperatures of NMRs isolated from the colony were also significantly higher than the ambient temperature. We also show that queens, but not subordinates, induce BAT thermogenesis in isolated, non-cold conditions. Our research provides novel insights into the role and mechanism of thermoregulation in this unique poikilothermic mammal.

## Introduction

Non-shivering thermogenesis in brown adipose tissue (BAT) helps maintain the body temperatures of homeothermic mammals in cold environments ^1^. BAT specifically expresses uncoupling protein 1 (UCP1), which dissipates the energy produced from lipid and glucose metabolism as heat, rather than adenosine triphosphate synthesis, by increasing the proton conductance of the inner mitochondrial membrane. Recently, BAT has also been shown to be involved in thermogenesis in non-cold environments in response to stimuli, such as diet, social defeat, handling, or the presence of an intruder ^2–5^. The detailed mechanisms involved in this process have been intensively explored to develop novel treatments for diabetes and other metabolic diseases, because BAT thermogenesis improves lipid and glucose metabolism ^6–9^. Interestingly, BAT has also been found in non-homeothermic mammals, which cannot maintain their body temperatures in cold environments ^10,11^. However, its function in these animals remains largely obscure.

The naked mole-rat (NMR; *Heterocephalus glaber*; Fig. 1a) is an African poikilothermic mammal that is hairless with thin skin and is known as the longest-living rodent in the world, with extraordinary cancer resistance ^12–14^. NMRs live in colonies comprising many individuals (average 70–80 individuals) and form complex underground systems of tunnels, which can reach a total length of 3–5 km per colony, and maintain a warm and relatively constant air temperature ^15,16^. These tunnels connect chambers that are used for different activities, including nests, toilets, food storage sites, and garbage spots. Interestingly, poikilothermic NMRs in a colony can temporarily maintain “behaviorally homeothermic” states, regulating their body temperatures by returning to their nest chamber and huddling together to share heat and become warmer, or moving to cooler areas within the tunnel network to decrease their body temperatures ^17,18^. In our laboratory, NMRs are housed in acrylic chambers connected by acrylic tunnels that are maintained at 30 ± 0.5°C, which represents a non-cold environment for NMRs (Fig. S1) ^19^. NMRs are also known for their unique eusociality – in a colony of up to 300 individuals, only one female (queen) and one to three males (kings) are reproductive, with all other members being sexually immature and working as subordinates ^20,21^.

**Figure 1.**
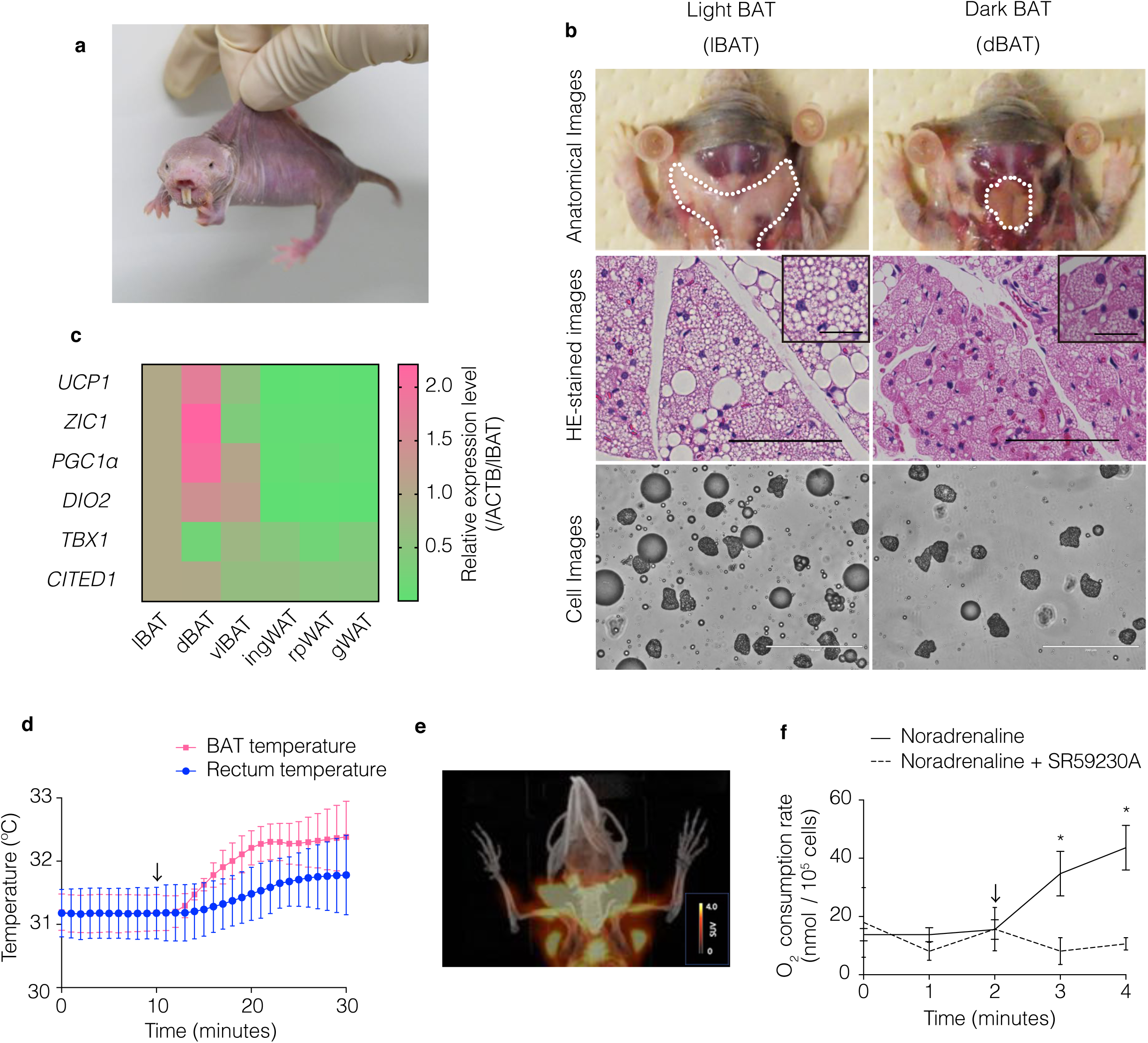
Poikilothermic naked mole-rats (NMRs; *Heterocephalus glaber*) possess thermogenic brown adipose tissue (BAT) **(a)** Photograph of an adult NMR. **(b)** Anatomical (top), hematoxylin–eosin (HE)-stained (middle) and cell (bottom) images of light BAT (lBAT) and dark BAT (dBAT). Scale bar = 100 µm for HE-stained images, 25 µm for insets, 200 µm for cell images. **(c)** Relative mRNA expression levels of brown (*UCP1, ZIC1, PGC1α* and *DIO2*) and beige (*TBX1* and *CITED1*) adipocyte marker genes reported in mouse and human in NMR adipose tissues. Expression levels were quantified by quantitative reverse transcription-polymerase chain reaction (qRT-PCR), normalized to beta-actin (*ACTB*), and compared with lBAT (*n* = 3 animals). vlBAT, ventral light BAT; ingWAT, inguinal white adipose tissue (WAT); rpWAT, retroperitoneal WAT; gWAT, gonadal WAT. **(d)** BAT and rectum temperatures of anesthetized NMRs before and after the *i*.*p*. injection of 1 mg/kg noradrenaline (arrow) at 30°C (*n* = 3 animals). **(e)** Positron emission tomography/computed tomography (PET-CT) imaging of NMR-BAT after the injection of 1 mg/kg noradrenaline and 11 MBq 2-deoxy-2-[^18^F]fluoro-D-glucose ([^18^F]FDG) at 32°C. **(f)** *In vitro* oxygen consumption rates of isolated adipocytes after the injection of 1 µM noradrenaline (arrow) with or without pre-incubation with 10 µM SR59230A (*n* = 3 animals per treatment), * *p* < 0.05 significantly different from SR59230A treated cells (paired *t*-test). All data are presented as means ± SEM with the exception of **(d)**, which are means ± SD.

Daly et al. have previously reported that NMRs have BAT in the interscapular region and in the area around the cervix ^10^. However, a previous work examining the thermogenic potential of NMR-BAT remains controversial. One previous study suggested that NMR-BAT can induce non-shivering thermogenesis based on indirect evidence from the *in vivo* oxygen consumption rate of NMRs after noradrenaline injection ^22^. However, to our knowledge, no direct evidence of NMR-BAT thermogenesis has been reported to date. Moreover, Buffenstein et al. have reported that NMRs cannot maintain their body temperatures in cold environments due to their inability to induce persistent non-shivering thermogenesis ^19^. NMR-specific mutations in the *UCP1* gene are thought to contribute to this inability ^23^. In general, the physiological role of BAT in poikilothermic NMRs is still completely unclear.

In this study, we investigated the molecular and histological characteristics of NMR-BAT, its thermogenic ability, and the NMR-BAT thermogenesis in physiological conditions. We demonstrate that NMRs possess a substantial amount of BAT with thermogenic activity. Although NMR-BAT thermogenesis slightly delays the decrease in body temperature in a cold environment, it was insufficient to maintain the body temperatures of the NMRs. Interestingly, NMR queens, but not subordinates, induce BAT thermogenesis when in an isolated, non-cold situation. Our results reveal that the NMR-BAT is indeed thermogenic and induces thermogenesis in physiological, cold, and non-cold environments, providing new insights into the process of thermoregulation in this poikilothermic rodent.

## Results

### Identification and examination of thermogenic BAT in naked mole-rats

We first performed a detailed characterization of NMR-BAT through dissection and a histological analysis. We identified two types of BAT around the cervix of NMRs: subcutaneous light BAT (lBAT), located in the interscapular region and around the cervix, and dark BAT (dBAT), located in deep regions under the cervical muscle (Figs. 1b and S2a). Hematoxylin–eosin (HE) staining and the isolation of adipocytes showed that dBAT mostly consisted of multilocular adipocytes, whereas lBAT was comprised of a mixture of multilocular and unilocular adipocytes (Figs. 1b and S2b). Because thermogenically active adipocytes contain smaller lipid droplets than inactive adipocytes ^24^, we measured the size of the lipid droplets in the HE-stained images, which showed that dBAT contained significantly smaller lipid droplets than the interscapular BAT of Crl:CD1 (ICR) mice (Fig. S2c). The average percentage of the total BAT per gram body weight was 1.88% (Fig. S2d).

Quantitative reverse transcription-polymerase chain reaction (qRT-PCR) showed that brown adipocyte marker genes reported in humans ^25^ and mice ^26^, such as *UCP1*, Zic family member 1 (*ZIC1*), peroxisome proliferator-activated receptor gamma coactivator 1 alpha (*PGC1α*), and iodothyronine deiodinase 2 (*DIO2*), were highly expressed in dBAT and lBAT (Figs. 1c and S2e). In contrast, for beige adipocyte, an inducible type of thermogenic adipocyte ^25, 26^, such as T-box transcription factor (*TBX1*) and transcriptional coactivator of the p300/CBP-mediated transcription complex (*CITED1*), were not upregulated in dBAT or lBAT (Figs. 1c and S2e). A gene ontology enrichment analysis further showed that processes related to the generation of precursor metabolites and energy, including the monocarboxylic acid metabolic process, acyl-CoA metabolic process, mitochondrial electron transport, electron transport from ubiquinol to cytochrome c, triglyceride metabolic process, response to fatty acid, glucose 6-phosphate metabolic process, and electron transport from cytochrome c to oxygen, were activated in dBAT, indicating that active metabolism occurs in this tissue (Fig. S2f). Furthermore, the western blotting showed that the UCP1 protein is highly expressed in dBAT and lBAT (Fig. S3).

To directly evaluate the thermogenic ability of NMR-BAT, we measured its temperature and the rectum temperature following the administration of noradrenaline by inserting a thermo-probe into the BAT and rectum of anesthetized NMRs. We found that noradrenaline injection caused the BAT temperature to increase by approximately 1.2°C, following which the rectum temperature gradually increased by 0.6°C (Fig. 1d). A positron emission tomography/computed tomography (PET/CT) analysis further showed that 2-deoxy-2-[^18^F]fluoro-D-glucose ([^18^F]FDG) was strongly taken up by the BAT of the noradrenaline-injected NMRs, with a clear “neck warmer”-like distribution of [^18^F]FDG around the cervix, in addition to its presence in the interscapular regions (Fig. 1e and Video S1).

To evaluate whether NMR-BAT thermogenesis depends on the adrenergic beta-3 receptor (ADRB3), which plays a critical role in BAT thermogenesis and has a relatively specific expression in the adipose tissues^1^, we measured noradrenaline-induced oxygen consumption rate of brown adipocytes isolated from a mixture of dBAT and lBAT after a noradrenaline treatment in the presence or absence of the ADRB3 inhibitor SR59230A, using adipocytes isolated from a mixture of dBAT and lBAT. We found that the stimulation with noradrenaline caused the rapid increase in oxygen consumption rate of brown adipocytes, but this increase was inhibited by the pre-treatment with SR59230A (Fig. 1f). Together, these findings indicate that poikilothermic NMRs possess ADRB3-dependent thermogenic BAT.

### Induction of NMR-BAT thermogenesis that slightly delays the decrease in body temperature in a cold environment

Next, we investigated the roles of NMR-BAT in physiological conditions. Because NMR skin is almost hairless and quite thin (Fig. 1a), BAT thermogenesis can be monitored by measuring the cervix surface temperature with a thermal camera. We found that the cervix surface temperature was correlated with the BAT temperature, as measured by the thermo-probe (Fig. S4). We measured the cervix temperatures of each NMR in the isolated situation because NMRs are known to enter a behaviorally homeothermic state by engaging in a heat-sharing behavior among others in their colony ^17,18^.

To evaluate the thermogenic ability of BAT in a cold environment for NMRs (20°C), we monitored the cervix surface temperature and the abdominal core body temperature of free-moving NMR subordinates using a thermal camera and a temperature telemetry system simultaneously. When NMRs were isolated from the colony and transferred to a room at 20 °C, the body and cervix surface temperatures gradually reduced, indicating NMRs were unable to keep their body temperature in cold environment as previously reported ^19^. We found that SR59230A initially accelerated the drops in both temperatures; however, SR59230A did not induce further decrease at equilibrium (Fig. 2a–c). These results suggest that, although BAT thermogenesis contributed to the delay in the decrease in body temperature after cold exposure, the NMRs were unable to maintain their body temperature at 20°C as previously reported ^19^ (Fig. 2a–c).

**Figure 2.**
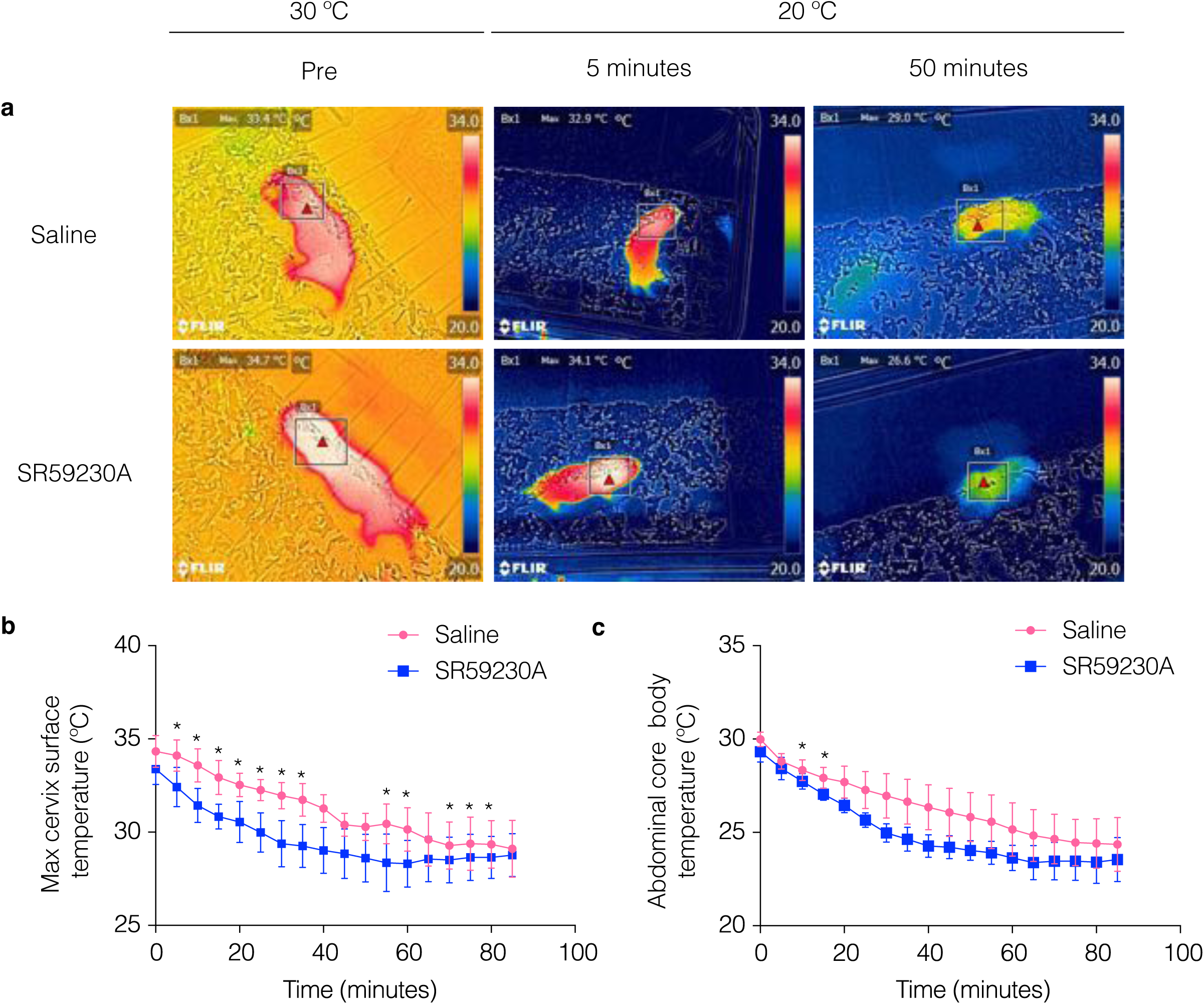
Brown adipose tissue (BAT) thermogenesis in naked mole-rat (NMR; *Heterocephalus glaber*) subordinates after cold exposure. **(a)** Thermal images of the declining body surface temperatures of NMRs during cold exposure (20°C) following the injection of saline or 20 mg/kg SR59230A (*n* = 4 animals per treatment). **(b)** Maximum cervix surface temperatures monitored by a thermal camera (*n* = 4 animals) and **(c)** abdominal core body temperatures recorded by a telemetry probe inserted into the abdominal cavity (*n* = 3 animals) of NMRs during cold exposure (20°C) following the injection of saline or 20 mg/kg SR59230A * *p* < 0.05 significantly different from SR59230A treated sample (paired *t*-test).

### Induction of BAT thermogenesis in naked mole-rat queens under isolated, non-cold conditions

We found that BAT thermogenesis had a slight but insufficient effect on supporting the body temperature of NMRs in a cold environment (Fig. 2). Therefore, we hypothesized that NMR-BAT may also play a role in non-cold environments. Interestingly, we found that NMRs isolated from their colony did not decrease their body temperatures and also showed higher body temperature than the ambient temperature (Fig. 3a). To test whether the thermogenesis of the isolated subordinates depended on BAT, we injected NMRs with the ADRB3 inhibitor SR59230A and again measured the body temperature of individuals after isolation. However, no significant change was observed in the subordinates (Fig. 3b, c).

**Figure 3.**
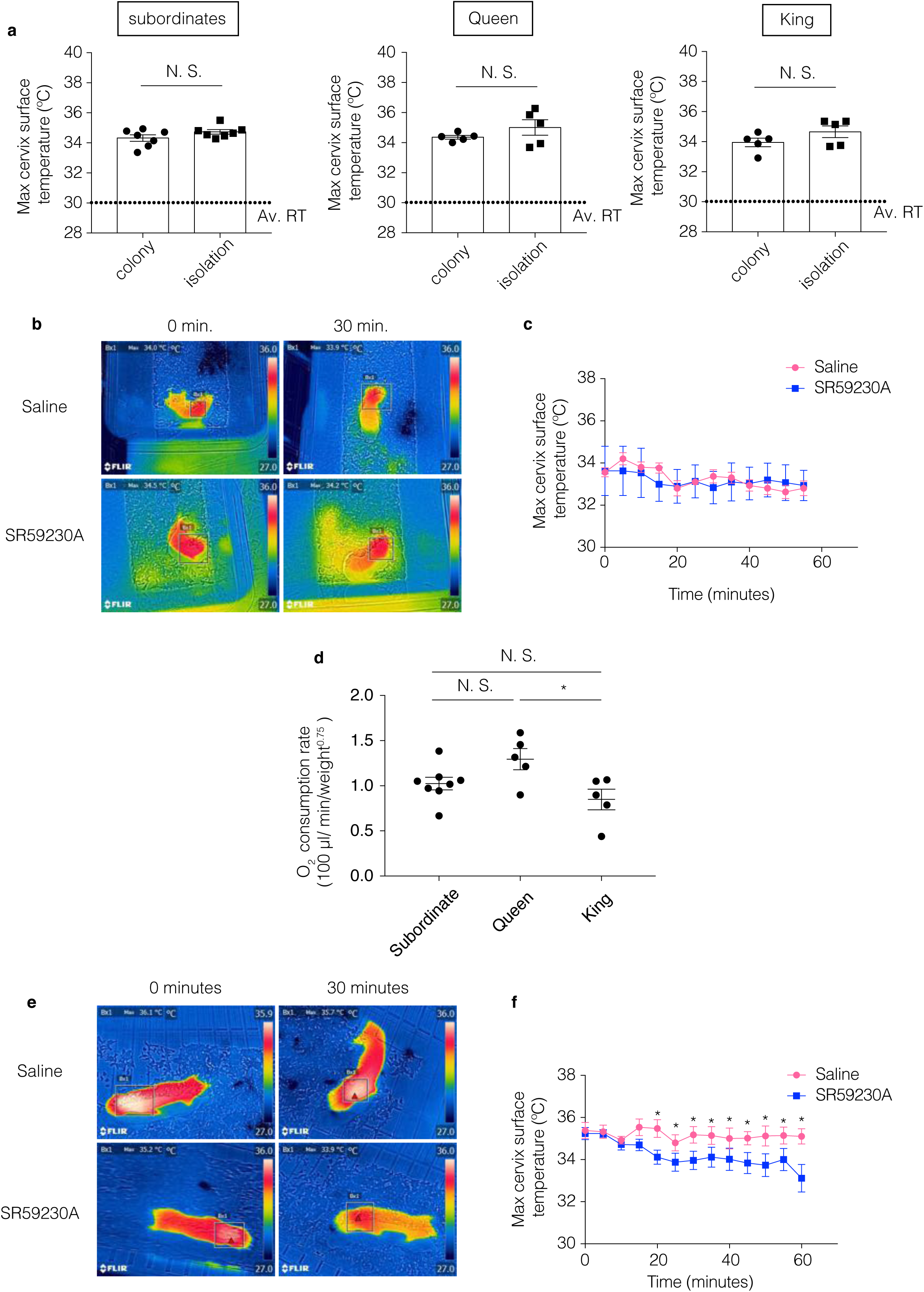
Isolated naked mole-rat (NMR; *Heterocephalus glaber*) queen exhibits BAT thermogenesis. **(a)** Maximum cervix surface temperatures of NMRs staying together in the colony (colony) or isolated from the colony (isolation). In the colony data, each point represents the average temperatures of individual NMRs measured at six times in the colony (three times in the nest and three times outside the nest). In the isolation data, each point represents the average temperatures of individual NMRs recorded every 30 min over 8 h. *n* = 5 animals for the queen and king, *n* = 7 animals for the subordinate. Av RT., the average room temperature, **(b)** Representative images and **(c)** time course of changes in the maximum cervix surface temperatures of socially isolated subordinates following the *i*.*p*. injection of saline or 20 mg/kg SR59230A (*n* = 3 animals). Measurement began after the isolation-induced increase in cervix surface temperature became stable [* *p* < 0.05 significantly different from SR59230A treated animals (paired *t*-test)]. Data are presented as means ± SEM. **(d)** Average oxygen consumption rates of the queen and subordinates in a socially isolated state measured in a metabolic cage in a non-cold environment (*n* = 5 queens, 7 subordinates, and 5 kings). Data collected over 4 h in the daytime after a 2-h habituation period. Data are presented as means ± SEM and were analyzed using one-way analysis of variance followed by Tukey’s multiple comparison test with a single pooled variance for multiple comparisons (* *p* < 0.05 significantly different from each groups), **(e)** representative images, and **(f)** time course of changes in the maximum cervix surface temperatures of socially isolated queens, following the *i*.*p*. injection of saline or 20 mg/kg SR59230A (*n* = 5 animals); * *p* < 0.05 significantly different from SR59230A treated animals (paired *t*-test). Data are presented as means ± SEM.

Notably, we found that the queens showed a tendency toward higher oxygen consumption rates than other members, suggesting that the BAT of queens may be more thermogenic than in other colony members in this situation (Fig. 3d). Although the body weight and age were higher in queen than in the subordinates ^27^, we also found that the body weight and age were weakly negatively correlated with the body temperatures of the isolated subordinate NMRs (Fig. S5a, b), suggesting that the higher thermogenic activity of queens was not related to their heavier body weights or older ages. No significant differences in body temperature were found between the sexes (Fig. S5c). To test whether the thermogenesis of the isolated queens depended on BAT, we injected NMRs with the ADRB3 inhibitor SR59230A and again measured the body temperature of individuals after isolation. As a result, we found that the body temperatures of the socially isolated queens significantly decreased following SR59230A injection (Fig. 3e, f). These results indicate that NMR queens, but not subordinates, activate BAT thermogenesis in an isolated, non-cold environment.

## Discussion

In this study, we provided direct evidence in support of the thermogenic potential of BAT in poikilothermic NMRs, which is ADRB3 dependent. We show that BAT thermogenesis was insufficient to maintain the body temperatures of the NMRs in a cold environment although it can slightly delay the decrease in the body temperatures of NMRs. Furthermore, in a non-cold environment, BAT thermogenesis contributes to maintaining the body temperature of the isolated queen. This research provides *in vitro* and *in vivo* evidence of NMR-BAT thermogenesis in physiological conditions that have not previously been studied.

A previous study suggested that the NMR *UCP1* gene has a unique sequence that may contribute to the inability of thermogenesis in NMRs ^23^; however, our results clearly show that, NMR-BAT does have thermogenic potential (although we did not compare the levels of thermogenic ability of NMR-BAT with those of other species). Whether NMR-BAT has more or less thermogenic potential than the BAT of other species is an important question for future research.

Although our results clearly showed the thermogenic ability of NMR-BAT, NMRs could not maintain their body temperatures in a cold environment (20 °C) as previously described^19^. This result may reflect the fact that NMRs have high heat dissipation because of their thin and hairless skin.

Interestingly, we observed that only queens induce BAT thermogenesis when in an isolated, non-cold situation. Previous reports in mice and rats have indicated that social isolation affects the metabolism and volume of adipose tissues, including BAT ^29,30^, and that psychological stresses, such as social defeat ^3^, handling, ^4^ or the presence of an intruder, ^5^ have been shown to induce BAT thermogenesis. Recent studies have revealed that cortisol concentration was also upregulated in NMRs after social isolation ^31,32^, suggesting that social isolation induces psychological stress in NMRs. Although the differences in stress levels between the queen and other colony members and the neurological systems to transmit the stress to BAT during social isolation in NMRs are still unknown, such differences may contribute to the observed difference in BAT thermogenesis in the isolated situation. Another possibility is that the difference in the BAT thermogenesis between the queen and the other members may result from differences in the thermogenic function or volume of BAT.

However, we were unable to collect and examine BAT from the queen by dissection due to the limited number of queens in our laboratory. Therefore, further experiments are required to investigate the mechanism underlying this queen-specific activation of BAT under isolation in a non-cold environment. Moreover, our observations suggest the possibility that the queen might be more resistant to decreasing body temperature in a cold environment than other members. However, we could not measure the core body temperature of the queen in a cold environment by inserting the thermo-probe into the abdominal cavity because this would have been too invasive, and the number of reproductive queens was quite limited. However, elucidating this factor will be another important issue for future works.

Moreover, in subordinates, we observed higher body temperatures than the ambient temperatures in a non-cold environment; however, the ADRB3 inhibitor did not decrease the body temperature. Although we cannot deny the possibility that more sensitive measurements can provide different results, our results may suggest that the isolated subordinates increase their body temperatures by BAT-independent mechanisms such as activity-dependent thermogenesis.

A current open question remains whether NMRs staying together in the colony also induce BAT thermogenesis. However, NMRs display the heat-sharing behavior only when inside the colony, and the intraperitoneal injection of SR59230A is not suitable for suppressing BAT thermogenesis for a long time period. Although we tried to irreversibly suppress BAT thermogenesis by cutting the sympathetic nerve projecting into the BAT, we failed because this method was too invasive. Developing methods that suppress BAT thermogenesis for long time periods and allow for a more sensitive measurement of BAT temperatures will contribute to the further understanding of the function and role of NMR-BAT.

In conclusion, we revealed that the poikilothermic NMR-BAT is thermogenic, inducing thermogenesis in physiological, cold, and non-cold environments. This work provides novel insights into the previously unclear role of BAT in this poikilothermic mammal. Further studies of BAT thermogenesis in the NMR and other non-homeothermic animals should continue to advance our understanding of the unexpected roles of BAT in animal homeostasis.

## Materials and Methods

### Study organisms

The NMRs used in this study were maintained at Kumamoto University, Kumamoto, Japan, where they were housed in four to 10 acrylic chambers that were connected by acrylic tunnels, at 30 ± 0.5°C and 55% ± 5% humidity with a 12 h light/12 h dark cycle (Fig. S1). The body temperatures of NMRs in the colony were assessed using 1- to 12-year-old NMRs. The effect of social isolation was assessed using 1- to 13-year-old subordinates, 5- to 13-year-old queens, and 6- to 12-year-old kings. JcI:ICR mice were purchased from CLEA Japan, Inc., and adipose tissues were collected from 1- to 2-year-old subordinates and 6-week-old mice for cytological and histological analyses.

All experimental procedures were permitted by the Institutional Animal Care and Usage Committees of Kumamoto University (Approval No. A30-043) and Hokkaido University (Approval No. 14-0065).

### HE staining and measurement of the lipid droplet size

The NMR and mouse adipose tissues were fixed with 4% paraformaldehyde in phosphate-buffered saline. HE staining was performed by Sapporo General Pathology Laboratory Co., Ltd. (Hokkaido, Japan), and images of the HE-stained samples were acquired with a BZ-X 710 fluorescence microscope (Keyence). The lipid droplet size was measured using a BZ-X image analyzer (Keyence).

### mRNA-sequencing analysis

RNA was extracted using Trizol reagent (Life Technologies) in accordance with the manufacturer’s protocol. A Qiagen RNeasy column was used for further purification, and genomic DNA was excluded using the TURBO DNA-free™ kit (Invitrogen). RNA quantity and quality were measured by Qubit (Invitrogen) and a 2100 Bioanalyzer using the RNA 6000 Nano Kit (Agilent Technologies). The TruSeq RNA Library Prep Kit v2 (Illumina) was used for library preparation in accordance with the manufacturer’s protocol. The acquired library was quantified using the High Sensitivity DNA Kit (Agilent Technologies) and Kapa Library Quantification Kit (Kapa Biosystems) with the Applied Biosystems ViiA7™ Real-Time PCR System (Applied Biosystems) using the manufacturer’s protocol. The library was loaded into a flow cell for cluster generation with the TruSeq Rapid SR Cluster Kit (Illumina) and was sequenced using the Illumina Hiseq 2500 System to obtain single-end 100-nucleotide sequences.

The NMR reference genome (HetGla_female_1.0) and annotation files downloaded from Ensembl 92 (https://www.ensembl.org) were used for data analysis. The acquired fastq files were trimmed using Trim Galore ver. 0.4.4. ^33^, and the transcript abundances (transcripts per million [TPM]) in the trimmed fastq files were calculated using RSEM ver. 1.2.31 with Bowtie 2 ^34,35^.

For the gene ontology enrichment analysis, the calculated TPM of dBAT was compared with that of inguinal white adipose tissue, and the top 200 upregulated genes in dBAT were analyzed by Metascape using gene ontology annotation of the mouse ^36^.

### qRT-PCR

Total RNA was extracted using the RNeasy Lipid Tissue Mini Kit (Qiagen), and genomic DNA was eliminated using the TURBO DNA-free Kit (Invitrogen) following the manufacturers’ protocols. Reverse transcription reactions were carried out using ReverTra Ace^®^ qPCR RT Master Mix (TOYOBO) with 300 ng of RNA as a template. The resulting cDNA was prepared for qPCR using Thunderbird^®^ qPCR Mix (TOYOBO) in a 384-well plate with the primers listed in Table S1. qPCR was performed on a CFX384 Touch™ Real-Time PCR Detection System (BIO-RAD).

### *In vivo* measurement of BAT thermogenesis

NMRs were anesthetized with 0.1 µg/g medetomidine hydrochloride (Dorbene^®^ Vet; Kyoritsu Seiyaku Co.), 4 µg/g midazolam (Dormicum^®^; Asteras Pharma Inc.), and 5 µg/g butorphanol (Vetorphale; Meiji Seika Pharma Co.). BAT and rectum temperatures were simultaneously measured using a thermo-probe (plastic-coated thermistor, 1 mm diameter). To measure the BAT, the skin above the interscapular region was incised without injury to the vasculature and nerves, and the thermo-probe was inserted under the lBAT. A thermo-probe was also inserted into the rectum at the same time. Once the BAT and rectum temperatures stabilized, 1 mg/kg noradrenaline was injected into the abdominal cavity, and the temperatures of the BAT and rectum were recorded over 30 min. All procedures were performed in a non-cold environment (30 ± 0.5°C), and the NMRs were placed on a hot plate at 32°C (NHP-M30N; NISSIN RIKA) during the experiment.

### *In vivo* measurement of oxygen consumption

The *in vivo* oxygen consumption rate was measured using an O_2_/CO_2_ metabolism-measuring system (MK-5000RQ6; Muromachi Kikai) and MMS-ML/6 software (Muromachi Kikai). The oxygen consumption of a single NMR was measured at 30 ± 0.5°C in a sealed chamber. The NMRs were habituated to the sealed chamber for 2 h prior to testing, and the oxygen consumption rate was recorded for 4 h during the daytime.

### *In vitro* measurement of adipocyte oxygen consumption

NMR brown adipocytes were isolated from lBAT and dBAT as previously described ^37^. Briefly, incised NMR adipose tissues were incubated in Krebs–Ringer bicarbonate-HEPES (KRBH) buffer (130 mM Na^+^, 4 mM K^+^, 0.75 mM Ca^2+^, 1 mM Mg^2+^, 121.5 mM Cl^−^, 10 mM HCO_3_^−^, 1 mM Mg^2+^, 4 mM HPO_4_^2−^, 30 mM HEPES, and pH 7.4) with 1% fatty acid-free bovine serum albumin (Wako), 6 mM glucose, and 1 mg/mL collagenase (Sigma) at 37°C for 1 h, with shaking at 90 rpm/min. After filtering the suspension through a 200 µM nylon filter, the filtrate was centrifuged at 50 × g for 2 min. The floating adipocytes were then collected and diluted with KRBH buffer containing 4% bovine serum albumin and 2.7 mM glucose and then recentrifuged and washed three times. The acquired adipocytes were incubated at room temperature for 1h before measurement.

The oxygen consumption rate was measured using a Clark-style oxygen electrode in a water-jacketed Perspex chamber at 37°C with the StrathKelvin 782 2-Channel Oxygen System (StrathKelvin Instruments). Once the stable oxygen consumption rate had been recorded, noradrenaline was injected into the chamber at a final concentration of 1 µM. To examine the effect of the noradrenaline receptor inhibition, SR59230A (10 µM; Sigma) was injected 10 min before injecting noradrenaline.

### Measurement of the body temperature in a cold environment via a thermal camera and telemetry probe

For the measurement of the cervical temperature and abdominal core temperature in a cold environment (20 °C), we first injected saline or 20 mg/kg SR59230A to the NMR abdominal cavity in a non-cold environment (30 °C). Then, we moved the NMR in the acrylic cage to a cold environment (20 °C) and measured changes in the cervical temperature with the thermal camera and the abdominal core temperature by telemetry probe every 5 min for 90 min. Body surface temperatures were monitored using a thermal camera (CPA-E6A; FLIR), and the acquired data were analyzed by FLIR Tools ver. 2.1 (FLIR). The abdominal core body temperature was measured using a telemetric probe (G2 E-mitter; STARR Life Sciences Corp.) and an ER4000 receiver (STARR Life Sciences Corp.). The G2 E-mitter was inserted into the abdominal cavity of anesthetized NMRs, and the animals were left for at least 7 days before being used in the experiment. The acquired data were analyzed by VitalView ver. 4.1 (STARR Life Sciences Corp.).

### Measurement of the body temperature via a thermal camera in a non-cold environment

For the measurement of the cervical temperature of the socially isolated NMRs, we measured the change in the cervical temperature via a thermal camera every 30 min for 8 hrs after 30 min social isolation. Individual NMRs were moved from the colony to the acrylic chamber at 30 ± 0.5°C and 55% ± 5% humidity with a 12 h light/12 h dark cycle. For the measurement of the body temperature in socially isolated NMR to compare against the body weight, age or sex, we measured the change in the cervical temperature via a thermal camera after 30 min of social isolation.

To measure the cervical temperature of the socially isolated NMR with the injection of saline or 20 mg/kg SR59230A to the NMR abdominal cavity, we measured the change in the cervical temperature using the thermal camera every 5 min for 90 min in a non-cold environment (30 °C) after 30 min of social isolation and the injection of saline or SR59230A.

### PET-CT imaging

For PET-CT imaging, NMRs were fasted overnight and kept at 30 ± 0.5°C and 60% humidity. The NMRs were administered 1 mg/kg noradrenaline, following which 11 MBq [^18^F]FDG was injected into the abdominal cavity. PET-CT images were then acquired 1 h after [^18^F]FDG administration using the Inveon small-animal multimodality PET/CT system (Siemens Medical Solutions). PET scanning was performed for 10 min followed by CT scan. During this experiment, the NMRs were maintained under isoflurane anesthesia and were kept at 32°C. Acquired PET images were reconstructed using the filtered backprojection algorithm with the ramp filter cut-off at the Nyquist frequency. The image matrix was 256 × 256 × 318, resulting in a voxel size of 0.388 × 0.388 × 0.398 mm^3^. CT images were reconstructed using a Feldkamp cone-beam algorithm, with a Shepp–Logan filter.

### Western blotting

Each adipose tissue was dissected, lysed in the buffer (125 mM Tris-HCl, pH 6.8; 4% SDS and 10% sucrose), and boiled for 10 min. After centrifuging, the supernatant was collected. The protein concentration was measured by TaKaRa BCA Protein Assay Kit (Takara Bio) in accordance with the manufacturer’s protocol. The 15 µg proteins were subjected to polyacrylamide gel electrophoresis, and then proteins were transferred to the polyvinylidene fluoride membrane. Blocking was performed with 0.5% skim milk for 1 h at room temperature. Blotted membranes were incubated with the primary antibody overnight at 4°C and the secondary antibody for 1 h at room temperature. We used anti-UCP-1 antibody (Sigma, U6382; 1:1000) and HRP-conjugated anti-rabbit IgG secondary antibodies (CST, #7074; 1:1000). The membrane was visualized by using Amersham ECL Prime Western Blotting Detection Reagent (GE Healthcare) and ImageQuant LAS 4000 Mini (GE Healthcare).

The detected membrane was washed by tris-buffered saline Tween-20 for 10 min. Then, the membrane was incubated with Ponceau-S staining solution for 15 min at a room temperature. After discarding the Ponceau-S staining solution, the membrane was incubated with 0.1% acetic acid for 2 min at room temperature. Then, the membrane was dried.

### Statistical analysis

GraphPad Prism (GraphPad) was used for the statistical analysis. Data were analysed using one-way analysis of variance followed by Tukey’s multiple comparison test with a single pooled variance for multiple comparisons or by Dunnet’s multiple comparison test with a single pooled variance. Two groups were compared using an unpaired *t*-test. All values are presented as means ± SD or means ± SEM, as noted.

## Acknowledgments

We thank Dr. T. Chujo for proofreading the manuscript; Prof. K. Seino, Prof. A. Takaoka, Prof. K. Tomizawa, Prof. Y. Ando, Dr. J. Kohyama, and professors at Institute for Genetic Medicine, Hokkaido University, Faculty of Life Sciences, Kumamoto University, and Institute of Molecular Embryology and Genetics, Kumamoto University for their administrative support and scientific discussion; Y. Sugiura for scientific discussion, Jane Doe of the Liaison Laboratory Research Promotion Center for technical support; I. Koya for the assistance of mRNA-sequencing; Y. Tanabe, Y. Fujimura, and M. Kobe for help with animal maintenance; T. Ohori for assistance in measuring NMR body surface temperatures; and all members of the K. Kimura and K.M. laboratories for technical assistance and scientific discussion. This research was supported in part by grants from the Ministry of Education, Culture, Sports, Science and Technology (MEXT) under grant number 26111006 (Grant-in-Aid for Scientific Research on Innovative Areas ‘Oxygen Biology: a new criterion for integrated understanding of life’), 15H05649 and 18H02365; PRESTO of the Japan Science and Technology Agency under grant number JPMJPR12M2; the Japan Agency of Medical Research and Development (AMED) under grant number JP19gm5010001.

K.M. was also supported by The Takeda Science Foundation, The Mitsubishi Foundation, The Naito Foundation, The Nagase Science and Technology Foundation, The Kurata Memorial Hitachi Science and Technology Foundation, The Nakajima Foundation, and The Suzuken Memorial Foundation.

## Contributions

Y.O. conducted most of the experiments; K.O., H.Y., K.H., and Y.K. conducted the PET-CT experiments; H.B., Y.K., S.M., A.S., and H.O. supported the mRNA-sequencing experiments and analysis; W.A., T.K., Y.S., and M.S. supported the characterization of brown adipose tissue; Y.O.-O., and K.K supported the BAT oxygen consumption rate measurements; Y.O., Y.O.-O., and K.M. designed the study; Y.O., Y.O.-O., and K.M. wrote the manuscript; Y.O.-O. and K.M. supervised the research.

## Accession code

RNA-seq data have been deposited in the DNA Data Bank of Japan database under accession code: DRA007737. Gene expression abundance data have also been deposited in the Genomic Expression Archive under accession code: E-GEAD-294.

---

**Figure S1.**
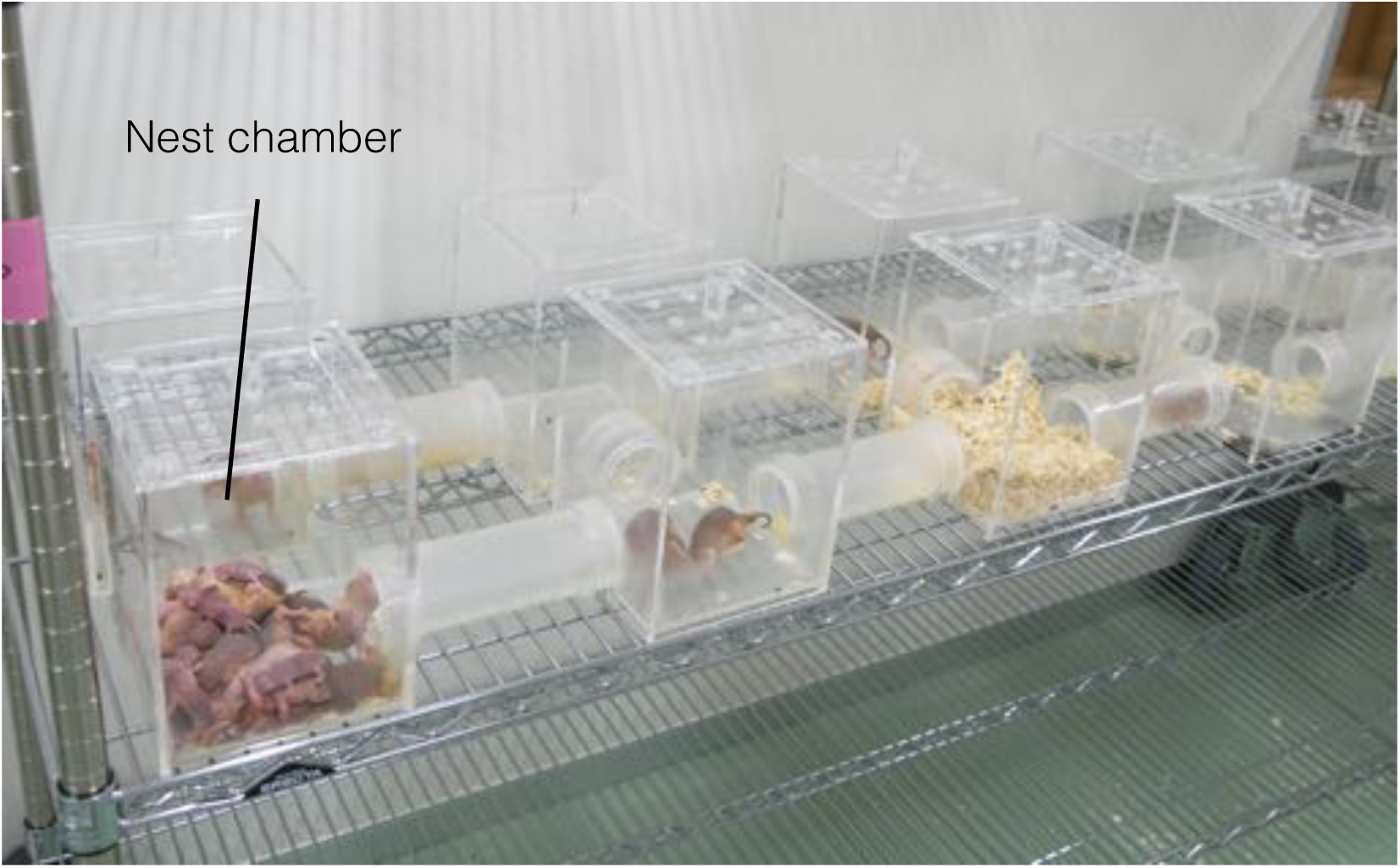
Images of acrylic chambers connected by acrylic tunnels used to house the naked mole-rats (NMRs; *Heterocephalus glaber*) in the laboratory. The NMRs were housed in acrylic chambers maintained at 30 ± 0.5°C, each of which was assigned a different use. In the nest chamber, NMRs shared heat and became warm by huddling together.

**Figure S2.**
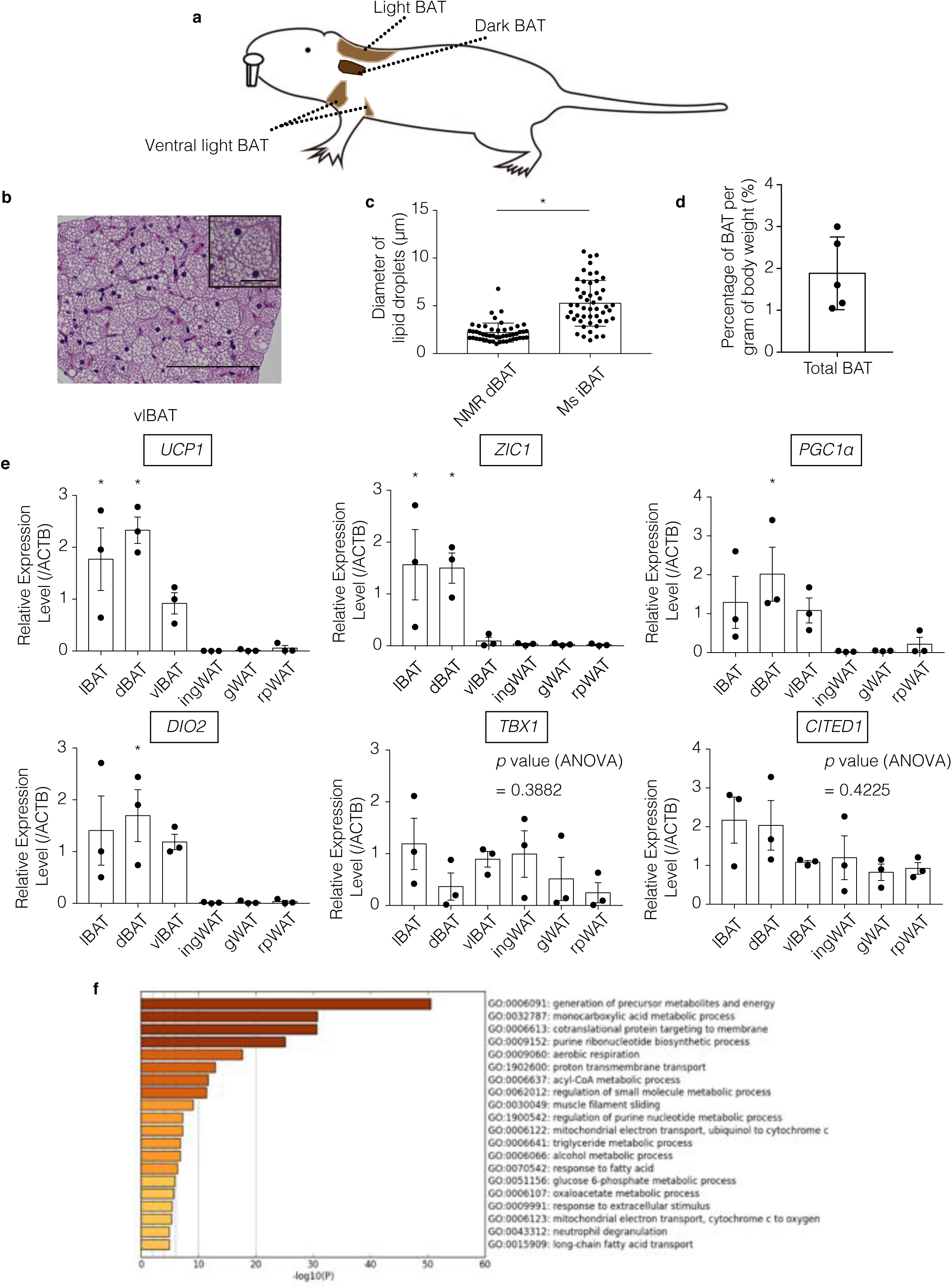
Characterization of naked mole-rat (NMR; *Heterocephalus glaber*) brown adipose tissue (BAT) **(a)** Schematic diagram showing the location of BAT in NMRs. **(b)** Hematoxylin–eosin (HE)-stained images of ventral light BAT (vlBAT). Scale bar = 100 µm for the HE-stained image, 25 µm for inset. **(c)** Lipid droplet size in NMR-dBAT and mouse-interscapular BAT (Ms iBAT). A total of 50 droplets were measured from three different fields of view per tissue type. The data were analyzed using an unpaired *t*-test. **(d)** Percentage of total BAT (light BAT [lBAT], dark BAT [dBAT], and vlBAT) per gram body weight. The data are presented as means ± SD (*n* = 5 animals). **(e)** Relative expression levels of brown or beige adipocyte marker genes reported in mouse and human in NMR adipose tissues. Expression levels were quantified by quantitative polymerase chain reaction (qPCR) using the primers listed in Table S1 and were normalized to beta-actin (*ACTB*) (*n* = 3 animals). ingWAT, inguinal white adipose tissue (WAT) (control); gWAT, gonadal WAT; rpWAT, retroperitoneal WAT. The data are presented as means ± SEM and were analyzed using one-way analysis of variance followed by Dunnett’s multiple comparison test with a single pooled variance (* *p* < 0.05 significantly different from ingWAT). **(f)** Gene Ontology enrichment analysis of the top 200 upregulated genes in the dBAT transcriptome using Metascape.

**Figure S3.**
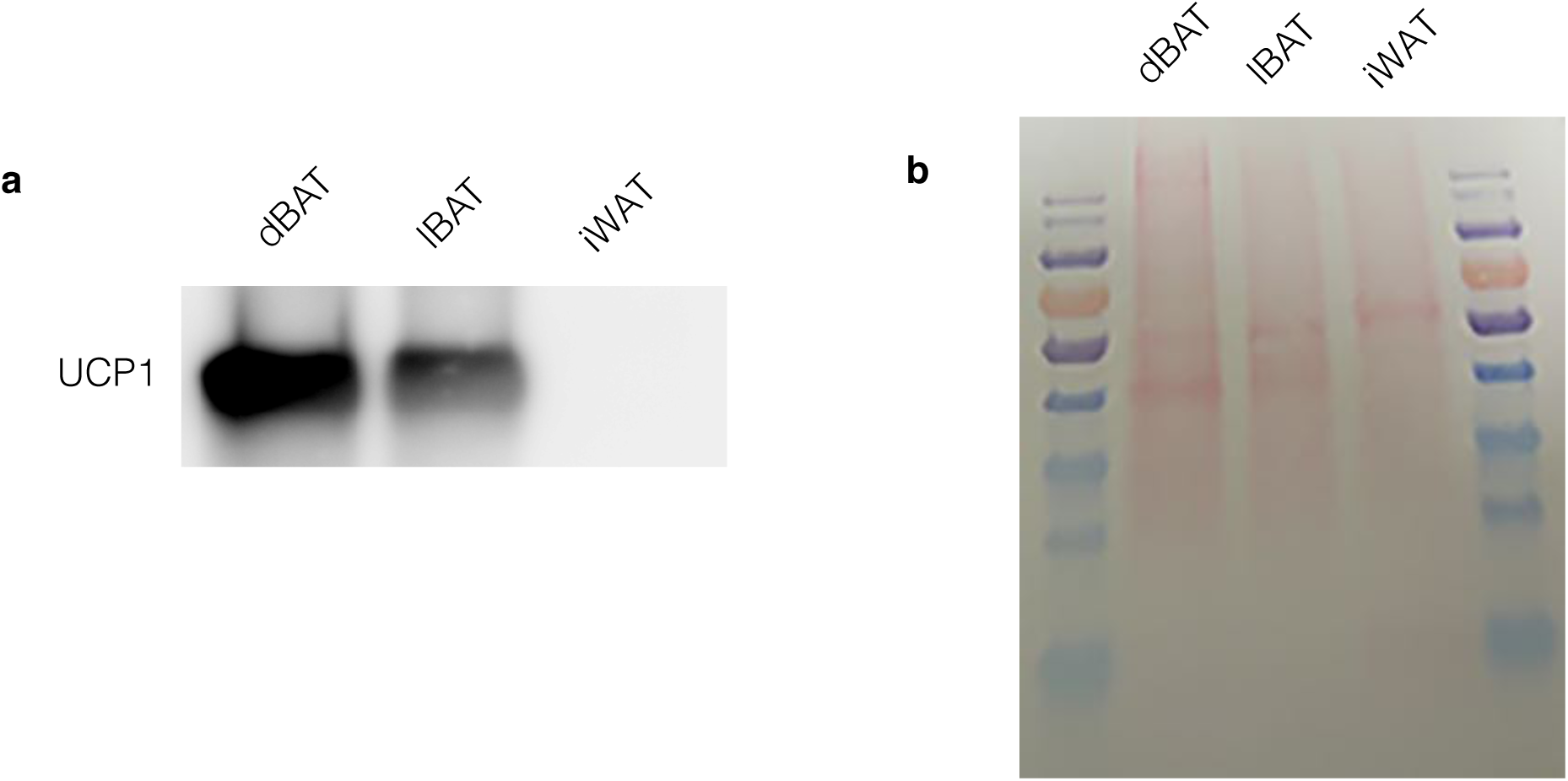
Uncoupling protein 1 (UCP1) expression in the naked mole-rat (NMR; *Heterocephalus glaber*) **(a)** UCP1 expression was evaluated by western blotting in NMR-dBAT, NMR-lBAT, and NMR-iWAT. **(b)** Ponceau-S staining was performed for the membrane used **(a)** after the UCP1 detection.

**Figure S4.**
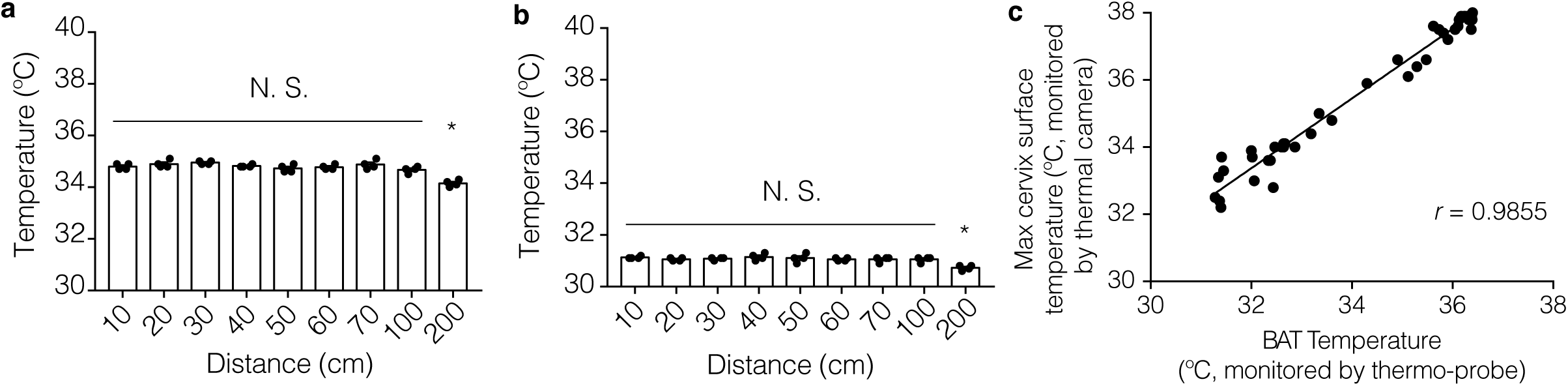
Correlation between brown adipose tissue (BAT) temperatures monitored with a thermo-probe and cervix surface temperatures monitored with a thermal camera. **(a, b)** Temperatures of the thermostable objects (**a;** 34.6°C–34.9°C, **b;** 31°C, measured by a mercury thermometer) were monitored by the thermal camera at various distances (four technical replicates at each point). Data are presented as means ± SEM * p < 0.05 (analyzed using one-way analysis of variance followed by Dunnett’s multiple comparison test with a single pooled variance (* *p* < 0.05 significantly different from 10 cm). In this experiment, we used the thermopack (Sugiyama-Gen, Japan) as a thermostable object. **(c)** BAT temperatures and the maximum cervix surface temperatures of anesthetized subordinates were simultaneously recorded by a thermo-probe and thermal camera, respectively, every 1 min during the increase in body temperature, following the injection of 1 mg/kg noradrenaline (*n* = 3 animals).

**Figure S5.**
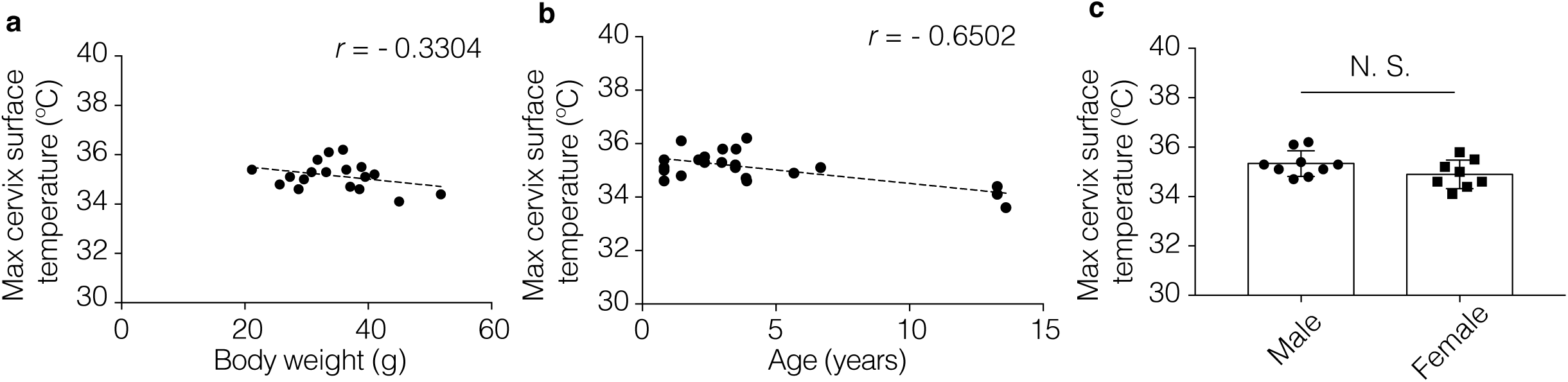
Relationships between the body temperature and body weight, age, and sex of socially isolated subordinate naked mole-rats (NMRs; *Heterocephalus glaber*) Maximum cervix surface temperatures of socially isolated subordinate NMRs monitored by thermal camera according to their (a) body weight, **(b)** age, and **(c)** sex. Data are presented as means ± SD. NS indicates nonsignificance (unpaired *t*-test).

**Video S1. Uptake of 2-deoxy-2-[18F]fluoro-D-glucose ([**^**18**^**F]FDG) in the brown adipose tissue (BAT) of naked mole-rats (NMRs; *Heterocephalus glaber*)**

The NMRs were administered 1 mg/kg noradrenaline, after which 11 MBq [^18^F]FDG was injected into the abdominal cavity. Positron emission tomography/computed tomography (PET-CT) images were then acquired 1 h after [^18^F]FDG administration. The oblong signal is a microchip for individual identification.

**Table S1.**
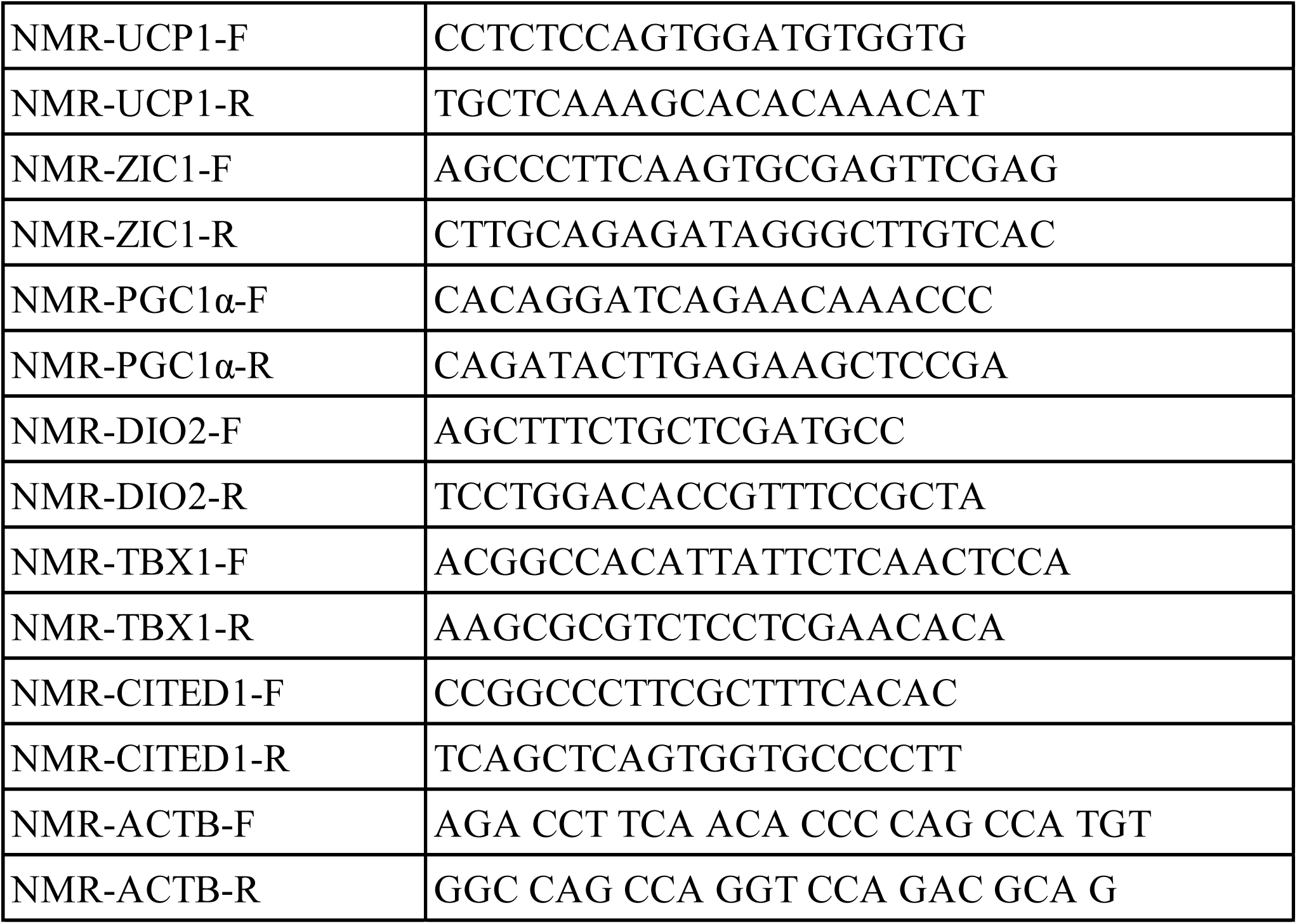
Primer Sequences.

